# CD239 deficiency in distal tubules impairs cell polarity and increases susceptibility to renal injury

**DOI:** 10.64898/2026.05.31.728029

**Authors:** Yamato Kikkawa, Jun Iwanami, Keisuke Hamada, Yuji Yamada, Takako Sasaki, Minoru Tanaka, Motoi Kanagawa

## Abstract

CD239, also known as Lutheran blood group glycoprotein (Lu) or basal cell adhesion molecule (B-CAM), is a transmembrane protein belonging to the immunoglobulin superfamily (IgSF). CD239 serves as a specific receptor for laminin α5, a subunit of laminin-511/-521, which are major components of renal basement membranes. A previous study of another group demonstrated that CD239-null mice are healthy and develop normally. Although no alteration in renal function was observed, most glomeruli in the mutant kidneys exhibited morphological abnormalities. In this study, we investigated the role of CD239 in renal tubules. We examined the distribution of CD239 using renal tubule-specific markers. CD239 was localized to the Henle’s loop, distal tubule, and collecting duct, but not to the proximal tubule. Next, we analyzed the localization of renal tubular molecules in CD239-null mice. The localization of uromodulin (UMOD) and Na–K–Cl cotransporter (NKCC2) was disrupted in the distal tubules lacking CD239, suggesting that CD239 plays a role in maintaining the polarity of renal epithelial cells. Furthermore, to examine the stability of the distal tubules, CD239-null mice were subjected to chronic kidney disease (CKD) using an adenine-rich diet. Blood analysis revealed that CD239-null mice fed an adenine diet readily developed CKD. Adenine-fed null mice exhibited more marked histological injury along the distal tubules compared to that by controls. These results indicate that CD239 is essential not only for maintaining cellular polarity but also for ensuring the stability of the distal tubules. Although CD239-null humans exhibit no obvious associated pathology, it could be a predisposition to CKD.

## INTRODUCTION

The kidneys play a crucial role in homeostasis by regulating water and electrolyte balance, vitamin metabolism, and hormone production (1). Homeostatic functions are mostly accomplished by the nephron epithelia of the glomeruli, proximal tubules, Henle’s loops, distal tubules, and collecting ducts (2). Renal epithelial cells must be highly polarized for proper renal function. Polarity is oriented in response to cell-cell and cell-matrix adhesions (3). All renal epithelial cells adhere to the basement membrane at the basal surface and to adjacent cells on the lateral side. The apical-basal polarity of epithelial cells results mainly from their adherence to the basement membrane via specific cellular receptors.

Renal basement membranes coat the entire outer surface of each individual nephron, collecting duct, and blood vessel. The basement membrane is a thin sheet of extracellular matrix composed primarily of laminin, Type IV collagen, nidogen, and sulfated proteoglycan (4). Among these, laminins play a pivotal role in the adhesion of epithelial cells to the basement membrane. Laminins are a family of non-collagenous glycoproteins composed of α, β, and γ chains. There are five α chains, three β chains, and three γ chains (5). Nineteen laminin isoforms have been identified in various cell and tissue cultures. Characterization of renal basement membranes has revealed that laminins provide molecular heterogeneity that corresponds in many ways to the segmental organization of the nephron (6). On the other hand, α5 chain, a subunit of laminin-511 and -521, is entirely contained in the renal basement membrane. Laminin α5 chain is bound by several different receptors, including integrin α3β1, α6β1, and α6β4 (7, 8), α-dystroglycan (9), and CD239 (10–12). Receptor binding also seems to contribute to the functional specificity manifested by distinct nephron segments.

Of the laminin receptors, CD239 specifically binds to laminin α5 (5, 12). CD239, also known as the Lutheran blood group glycoprotein (Lu) or basal cell adhesion molecule (BCAM), is a transmembrane protein belonging to the immunoglobulin superfamily (IgSF). Lu and BCAM have five disulfide-bonded extracellular IgSF domains (V-C1-I-I-I), one transmembrane domain, and a cytoplasmic tail (13, 14). Lu and BCAM are distinguished by their different cytoplasmic tails that arise from alternative RNA splicing. Lu is a spliceform with a longer cytoplasmic tail, whereas BCAM is another spliceform that has a cytoplasmic tail 40 amino acids shorter than Lu. Hereafter, when Lu and BCAM are indistinguishable, they are referred to as CD239. CD239 has mainly been studied as an antigen in the Lutheran blood group system and in the context of sickle cell disease (15). Furthermore, the receptor is overexpressed in various carcinoma (16–20). CD239 is expressed in most organs, including red blood cells (21, 22). To examine the roles of CD239, the gene was inactivated in mice (23). Although they were expected to exhibit disturbed morphogenesis and differentiation leading to lethality similar to laminin-α5 knockout mice (24), CD239-null mice were viable and fertile. A detailed analysis showed that the loss of CD239 causes thickening of the glomerular basement membrane (GBM) in the kidney and abnormal organization of smooth muscle layers in the intestine (23). We also generated CD239-null mice and showed that CD239 regulates biliary tissue remodeling in ductular reaction during the regeneration of the injured liver (25). Previous studies using CD239-null mice led us to hypothesize that the receptor plays a role in the kidney under loading conditions, leading to chronic renal disease.

In this study, we determined the distribution of CD239 in adult mouse kidneys using renal segment-specific markers. The kidneys of CD239-null mice also revealed that this receptor is associated with the maintenance of cellular polarity in the renal tubules. Furthermore, an experiment with an adenine-feeding load indicated that CD239 is involved in maintaining the stability of the distal renal tubule. Hence, our findings suggest that the deficiency of CD239 in the kidney influences the progression of renal dysfunction, such as in chronic kidney disease (CKD).

## RESULTS

### Distribution of CD239 in renal parenchyma

CD239 generates long- and short-cytoplasmic tail isoforms through alternative splicing. In mice, long-tail CD239 is expressed as a unique isoform in various tissues, including the heart, lungs, kidneys, intestines, and skeletal muscle (21, 26). To confirm the presence of the long-tail isoform in adult mouse kidneys, we performed immunohistochemistry using two antibodies that recognize the extracellular and cytoplasmic domains (Fig. 1a). Because the staining patterns of both antibodies merged, the longer form of CD239 was expressed in mouse kidneys. In contrast, the short-tailed form does not appear to be expressed in the kidney. We also reviewed the expression of CD239 in mouse kidneys by using sections cut in the transverse plane. The renal parenchyma is divided into two major structures: the renal cortex and medulla. The renal medulla is composed of the outer and inner medulla. The outer medulla is further divided into inner and outer stripes. CD239 was highly expressed in the inner stripes of the outer medulla and inner medulla, but not in the outer stripes of the outer medulla and cortex (Fig. 1b). In the renal cortex, as shown in previous studies, CD239 is expressed in the podocytes of the glomeruli (23, 26) (Fig. 1S and Table 1). Moreover, a subset of the renal tubules was CD239-positive in the outer stripe of the outer medulla and cortex. In blood vessels, the expression of CD239 was mainly observed in arterioles with smooth muscle layers but not in capillaries (Fig. 2S and Table 1).

**Figure 1.**
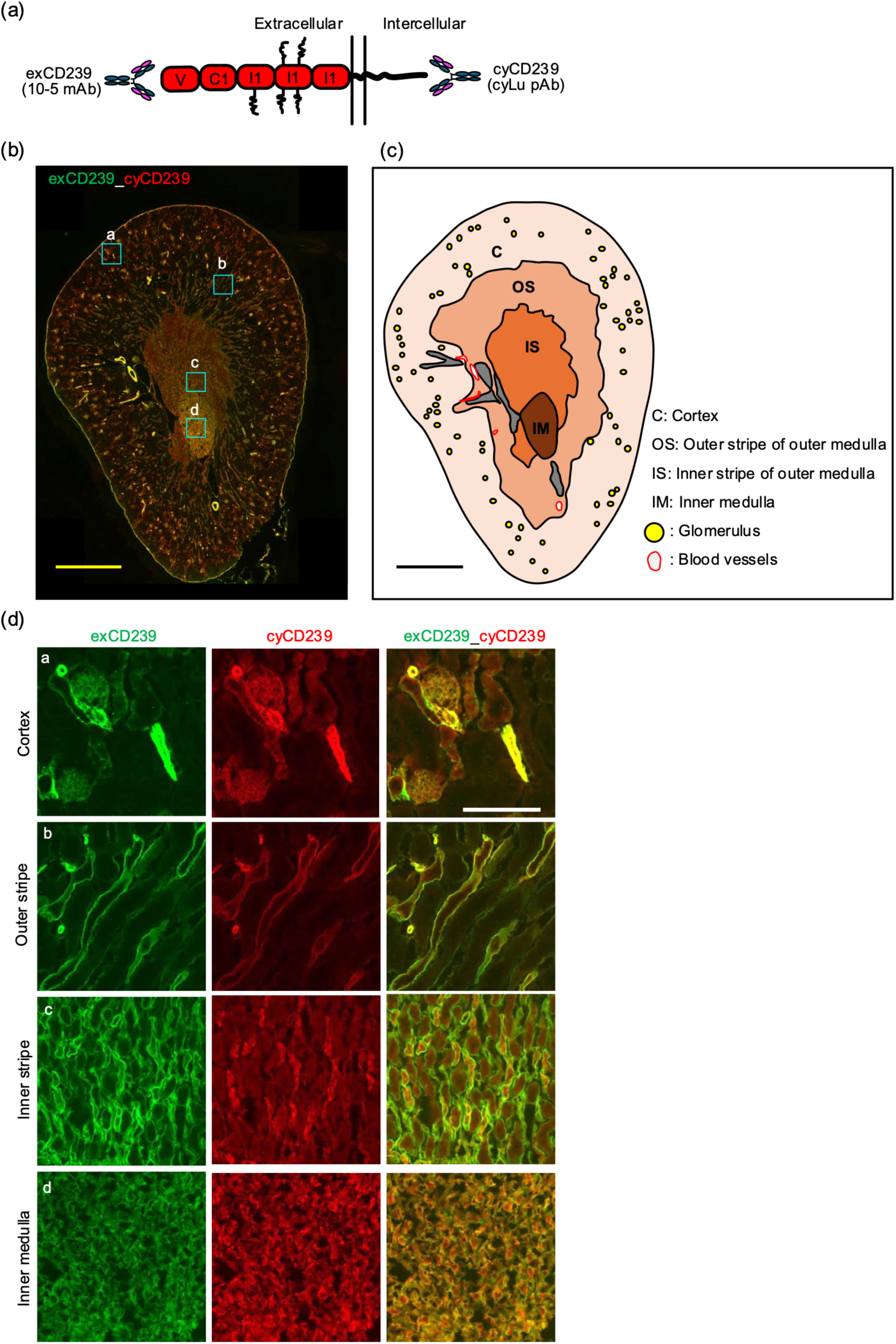
Distribution of CD239 in the adult mouse kidney. (a) Epitopes of two antibodies against CD239. (b) Immunostaining of transverse section through the adult mouse kidney. The sections were doubly stained with antibodies against extracellular (exCD239, green) and cytoplasmic (cyCD239, red) domains of CD239, respectively. Bar: 1.0 mm. (c) Schematic of the left transverse section of the adult mouse kidney and its anatomical regions. (d) High magnification images in each anatomical region. The locations are indicated by blue squares on the image of the whole kidney section. Bar: 100 μm.

**Figure 2.**
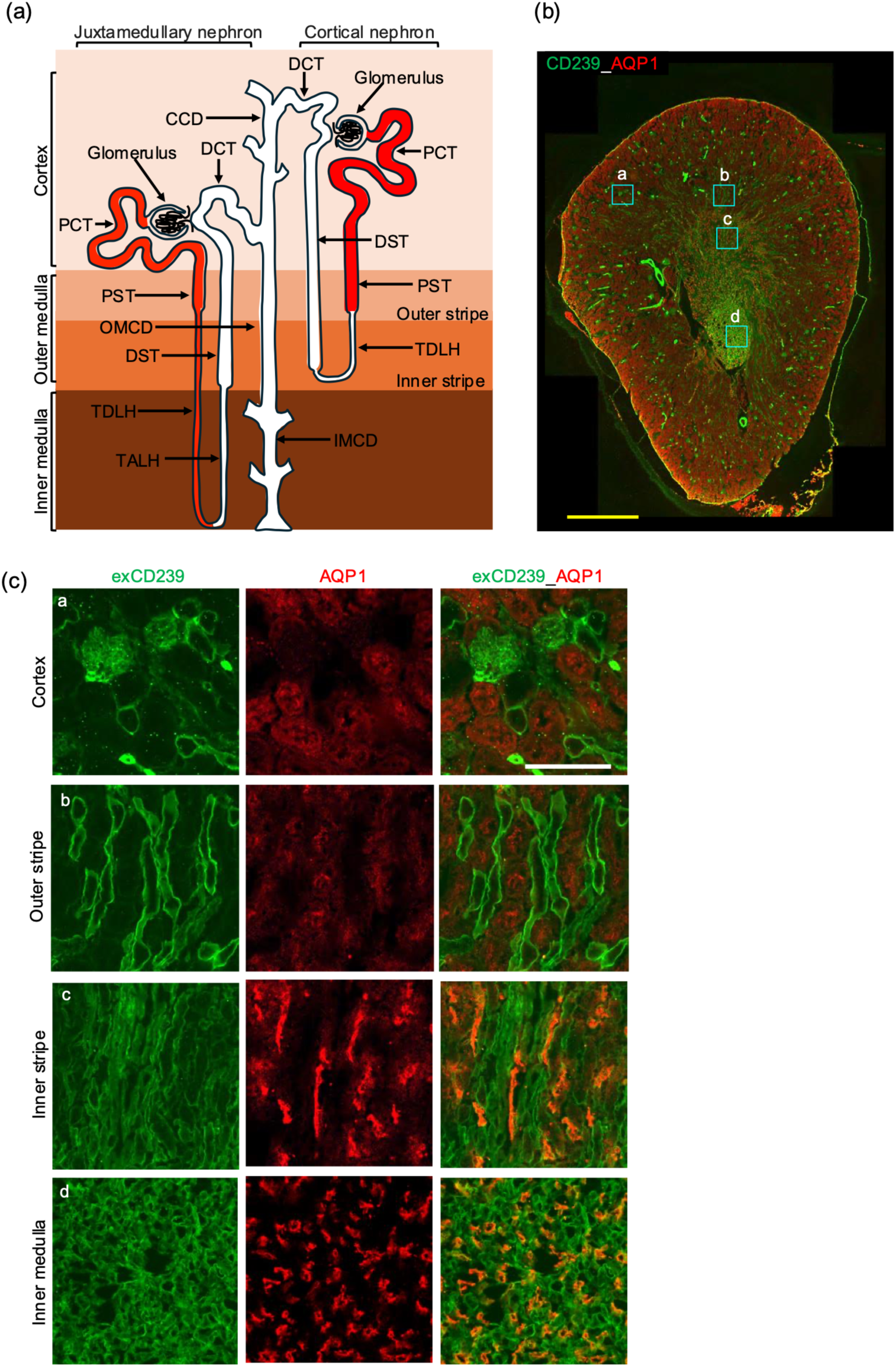
Expression of CD239 in Proximal convoluted tubule (PCT), proximal straight tubule (PST), and thin descending limb of Henle’s loop (TDLH). (a) Schematic of juxtamedullary and cortical nephrons. PCT, PST and TDLH are defined with localization of AQP1 (red). (b) Immunostaining of a whole mouse kidney section. The sections were doubly stained with antibodies against exCD239 (green) and AQP1 (red), respectively. Bar: 1.0 mm. (c) High magnification images in each anatomical region. The locations are indicated by blue squares on the image of the whole kidney section. Bar: 100 μm.

**Table 1.** Summary of the distribution of CD239.

| Table 1 Summary of the distribution of CD239 |  |  |  |  |  |  |  |  |  |  |  |  |  |  |  |  |
| --- | --- | --- | --- | --- | --- | --- | --- | --- | --- | --- | --- | --- | --- | --- | --- | --- |
|  | Glomerulus |  |  |  | PCT | PST | TDLH |  | TALH | DST | DCT | CD |  |  | Artery | Capillary |
|  | Podocytes | Glomerular capillary | Mesangial cells | Bowman's capsule |  |  | AQP1 positive | AQP1 negative |  |  |  | CCD | OMCD | IMCD |  |  |
| Tissue-specific marker | NPHS2 | CD146 |  |  | AQP1 | AQP1 | AQP1 |  | UMOD | UMOD | CALB1 | AQP2 | AQP2 | AQP2 | CD146 | CD146 |
| CD239 | ++ | - | - | - | - | - | ++ | + | ++ | +++ | + | ++ | ++ | ++ | +++ | - |
| Laminin $\alpha 5$ | +++ | | + | +++ | +++ | +++ | +++ | +++ | +++ | +++ | +++ | +++ | +++ | +++ | +++ | +++ |

PCT: proximal convoluted tubule, PST: proximal straight tubule, TDLH: thin descending limb of Henle's loop, TALH: thin ascending limb of Henle's loop, DST: distal straight tubule (synonym: thick ascending limb), DCT: distal convoluted tubule

### Localization of CD239 in nephron segments and collecting ducts

The adult mouse kidney is a complex organ containing more than 8000 mature nephrons connected to the collecting duct (27). Each nephron is subdivided into the renal corpuscle and several distinct tubular segments. Although nephrons are histologically complex, several antibodies can define renal corpuscles and tubular segments (28). To identify the renal tubular segments expressing CD239, we performed immunohistochemistry using antibodies against aquaporin 1 (AQP1), uromodulin (UMOD), and calbindin (CALB1), and aquaporin 2 (AQP2). AQP1 is localized in the proximal convoluted tubule (PCT), proximal straight tubule (PST), and thin descending limb of Henle’s loop (TDLH). CD239 was not expressed in PCT and PST, but was observed in TDLH (Fig. 2). UMOD is a marker of thin ascending limb of Henle’s loop (TALH) and distal straight tubule (DST). CD239 was distributed in the UMOD-positive renal tubules (Fig. 3). CD239 was weakly expressed in distal convolved tubule (DCT)-expressing CALB1 cells (Fig. 3S). Furthermore, collecting duct (CD) cells moderately expressed this receptor (Fig. 4S). Together with these results, CD239 was localized to Henle’s loop, distal tubule, and collecting duct, but not to the proximal tubule (Table 1).

**Figure 3.**
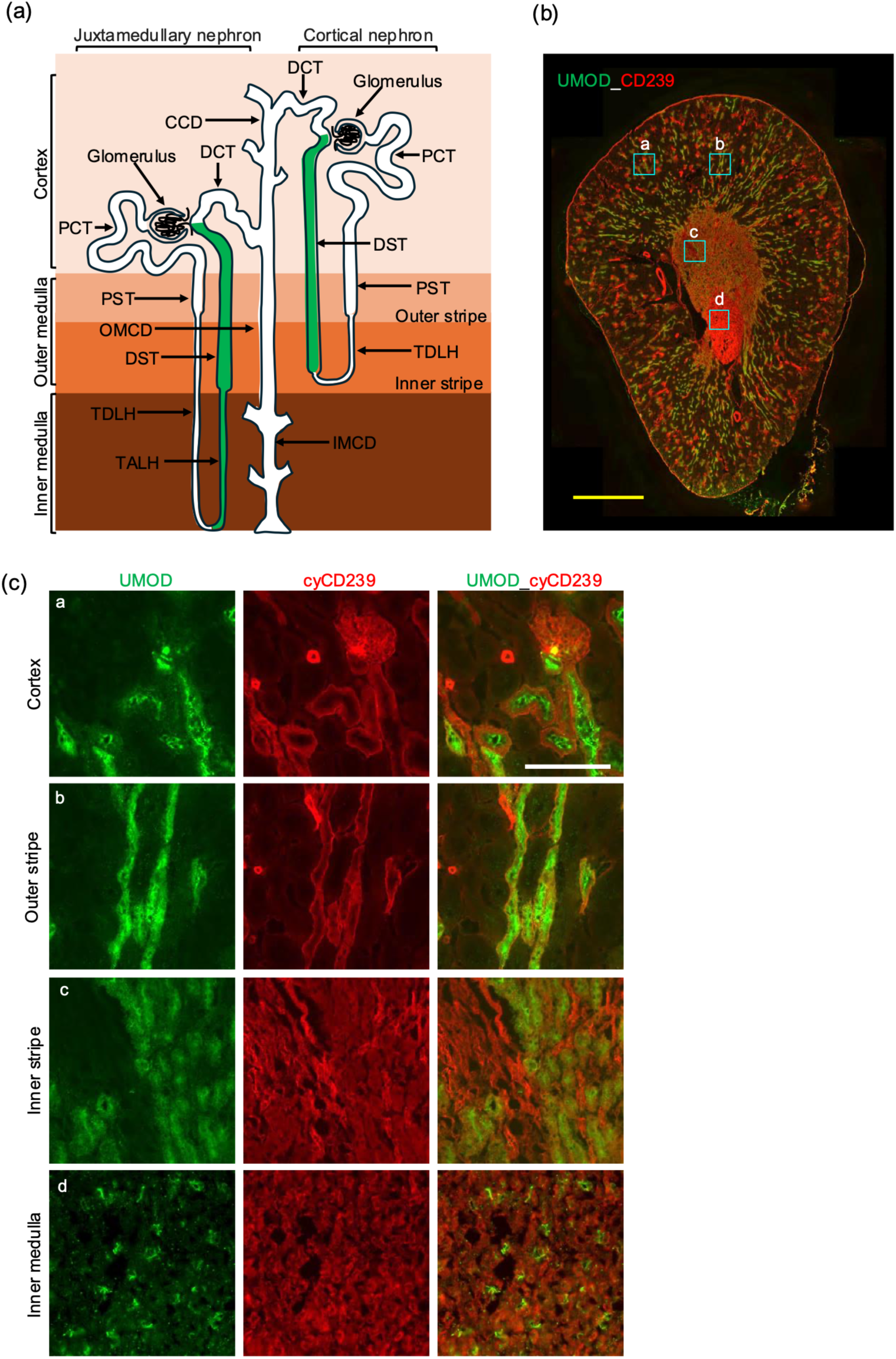
Expression of CD239 in thin ascending limb of Henle’s loop (TALH) and distal straight tubule (DST). (a) Schematic of the juxtamedullary and cortical nephrons. TALH and DST are defined with localization of uromodulin (UMOD, green). (b) Immunostaining of a whole mouse kidney section. The sections were doubly stained with antibodies against UMOD (green) and cyCD239 (red), respectively. Bar: 1.0 mm. (c) High magnification images in each anatomical region. The locations are indicated by blue squares on the image of the whole kidney section. Bar: 100 μm.

**Figure 4.**
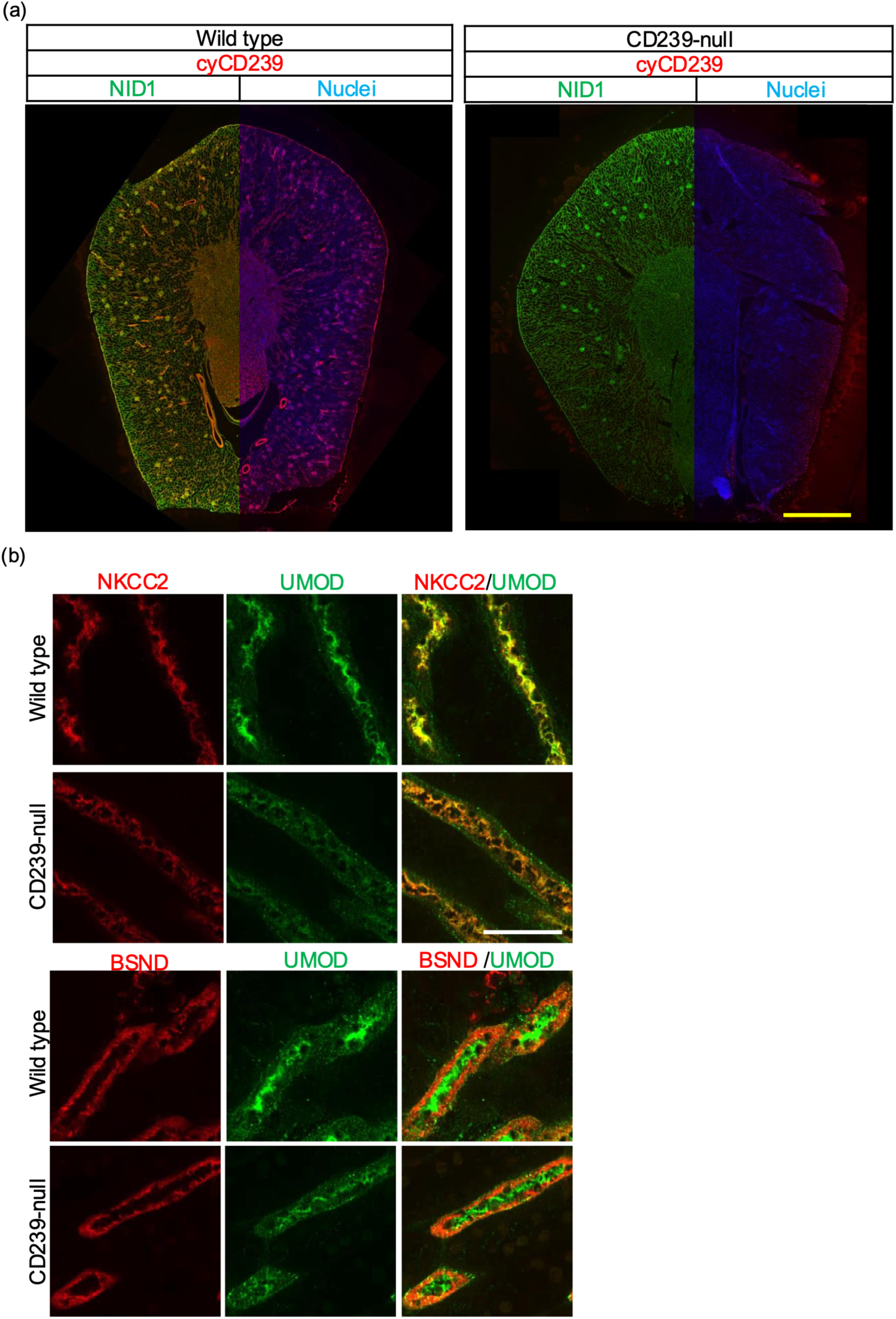
Localization of UMOD, Na-K-Cl cotransporter 2 (NKCC2) and Barttin (BSND) in DST of CD239-null mice. (a) Expression of CD239 in wild type and mutant kidneys (12-week-old female). The staining of nidogen-1 (NID1, green) was used to visualize basement membrane. Nuclei were counterstained with Hoechst 33258 (blue). (b) The kidney sections of wild type and CD239-null mice were doubly stained with antibodies indicated in the panels. DST in outer stripe of outer medulla was defined with morphology and UMOD staining. Bar: 50 μm.

### Cellular localization of renal tubular molecules in CD239-null mice

The long-tail isoform of CD239, Lutheran glycoprotein, has a dileucine motif in its cytoplasmic domain as a basolateral sorting signal (29). In support of this, our immunohistochemical analysis showed that CD239 was predominantly localized at the basal surface of renal epithelial cells. Although renal tubules lacking CD239 did not show any morphological abnormalities (Fig. 4a), we hypothesized that CD239 was associated with cellular polarity. Therefore, we explored the localization of renal tubular molecules involved in the metabolic function of the nephron segments and collecting ducts. In this study, we focused on three renal tubular molecules that are metabolically active in CD239 positive-renal tubules. Uromodulin (UMOD), also known as the Tamm-Horsfall protein, is the most abundant protein excreted in ordinary urine (30). UMOD is mainly located in the apical plasma membrane of the epithelium of the thick ascending limb. The localization of UMOD was disrupted in distal tubules lacking CD239 (Fig. 4b). NKCC2 is a cotransporter that reabsorbs approximately 20–25% of filtered NaCl in the thick ascending limb (31). This cotransporter is selectively expressed in the apical membrane of epithelial cells. The localization of NKCC2 in the apical plasma membrane is disordered in the distal tubules of CD239-null mice. Barttin (BSND) is a β subunit of CLC-chloride channels that localize to basolateral membranes of renal tubules (32). This subunit is required for the trafficking of CLC-chloride channels to the plasma membrane. The localization of BSND was not altered in the distal tubules lacking CD239.

### Stability of distal tubules in CD239-null mice

Although the localization of UMOD was disturbed in DST lacking CD239, alterations in renal function were not observed under normal conditions. The DST reabsorbs sodium and chloride from filtered urine. This segment also plays a role in potassium secretion and regulates the calcium, magnesium, and acid levels. Our previous study using CD239-null mice showed that CD239 influenced regeneration of injured liver (25). Therefore, we hypothesized that CD239 plays a role in the kidneys under loaded conditions, leading to chronic renal disease. To investigate the function of CD239 in DST, null mice were fed a diet supplemented with 0.2% adenine for 4 weeks. At physiological concentrations, adenine is metabolized to uric acid, which is excreted by the kidneys as xanthine. However, excess adenine is often metabolized through another pathway catalyzed by xanthine oxidase and converted from adenine to 2,8-dihydroxyadenine (DHA), which is poorly soluble at normal urine pH. Deposited DHA crystals injure renal tubules. In animal models, the excess feeding of adenine has been used to develop an experimental model of CKD in rodents (33, 34). After adenine feeding, although the body weights of the control and CD239-null mice decreased similarly, renal hypertrophy was observed in kidneys lacking CD239 (Fig. 5a). Blood samples were collected from mice for biochemical analyses. Blood urea nitrogen (BUN) and creatinine (CRE) levels significantly increased in CD239-null mice fed an adenine-supplemented diet (Fig. 5b). Alterations in total protein, albumin, Na, K, CL, Ca, Uric acid (UA), Inorganic phosphorus (IP), and Total cholesterol (T-CHO) levels were not observed in the plasma of CD239-null mice (Fig. 5S). Glucose (GLU) levels significantly decreased in adenine-fed CD239-null mice. Together, these results show that CD239 deficiency increases susceptibility to CKD.

**Figure 5.**
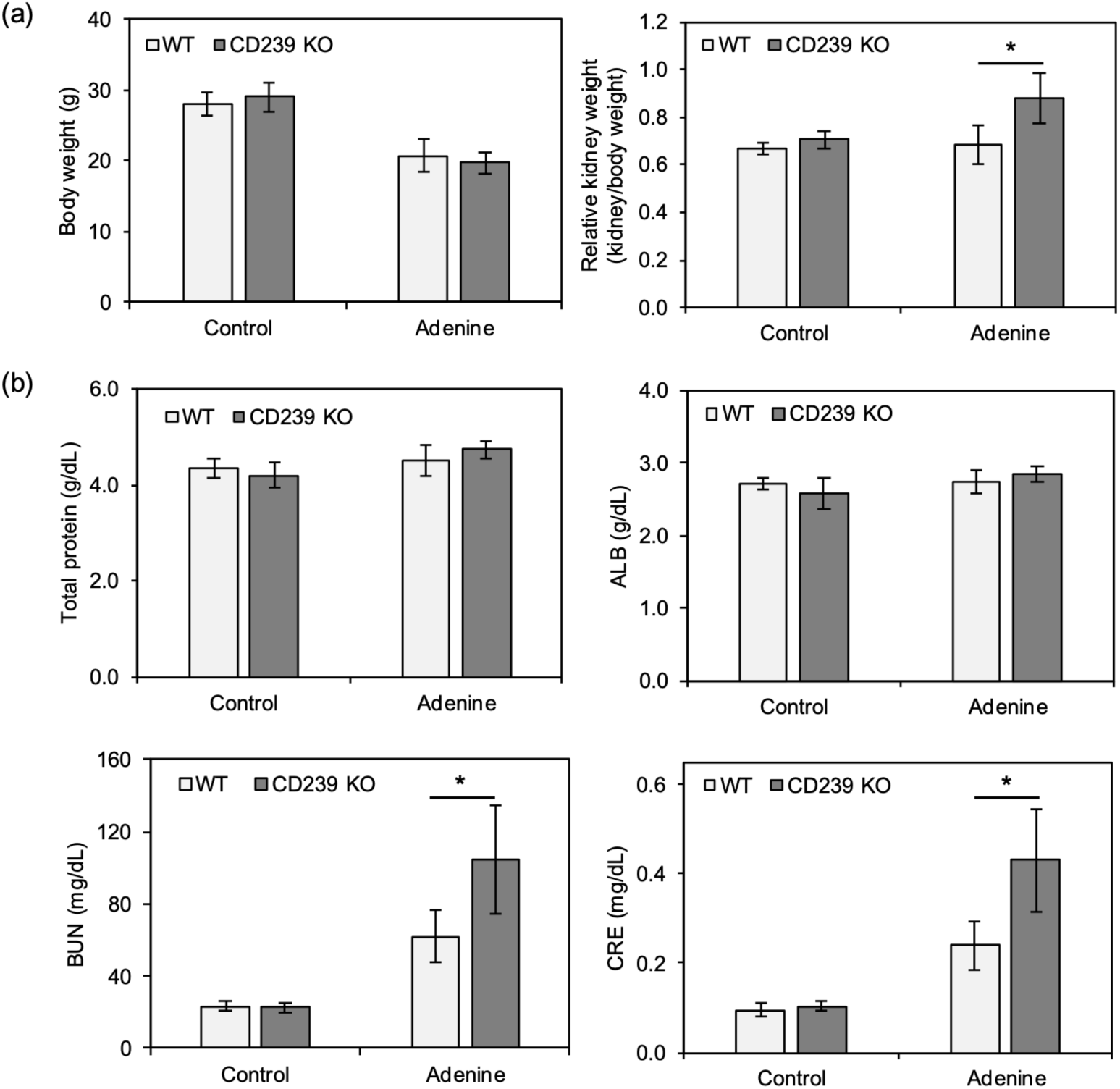
Blood biochemical analysis in the adenine-induced kidney injury mouse model. (a) Body weight and relative kidney weight of mice fed with adenine-supplemented diet. (b) The influences of adenine-feeding on serum markers of kidney function. After feeding adenine-supplemented diet, the blood was collected from wild type and CD239-null mice. The plasma was analyzed for total proteins, albumin (ALB), blood urea nitrogen (BUN), and creatinine (CRE). *, p < 0.05 by Student’s t-test.

To confirm the results of blood biochemical analysis, we performed histological analysis (Fig. 6). Macrophages mediate injury in adenine-induced CKD (35). Antibodies against F4/80 were used to label the macrophages. F4/80-positive macrophages proliferated and spread along the distal tubules in adenine-fed mice (Fig. 6). Moreover, CD239-null mice fed with the adenine diet showed more severe injury along the distal tubules than that in the controls. The results showed that CD239 is involved in maintaining the stability of the distal renal tubules.

**Figure 6.**
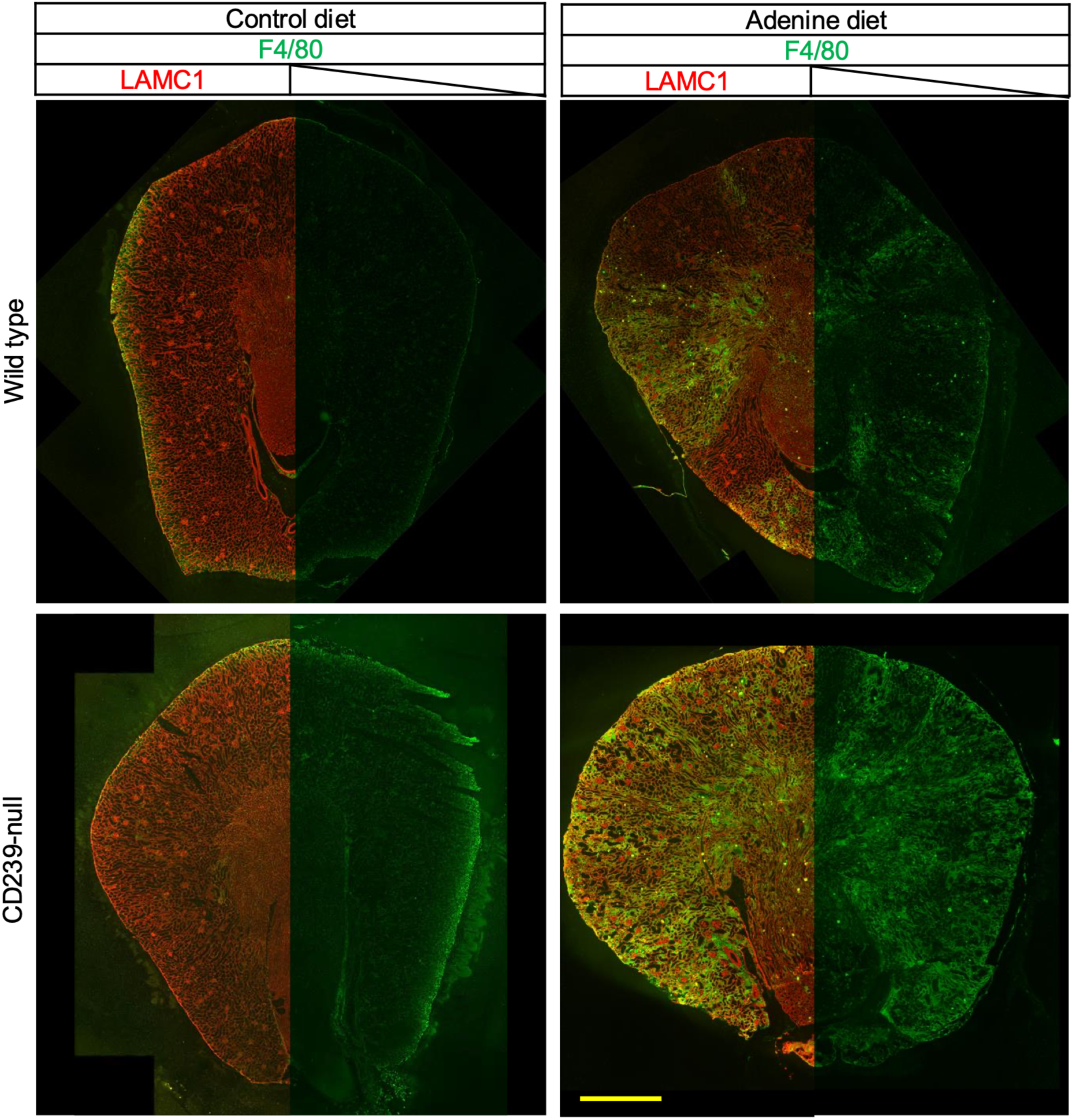
Immunohistological analysis in the adenine-induced kidney injury mouse model. Cryosections were prepared from the kidneys of wild type and CD239-null mice fed with a diet supplemented with 0.2% adenine for 2 weeks. The injured areas were defined with F4/80-positive macrophages, and the renal tubules were visualized with laminin γ1 (LAMC1) staining. They were doubly stained with antibodies against F4/80 (green) and LAMC1 (red). Bar: 1.0 mm.

## DISCUSSION

As described previously (21, 26), the long-tail isoform of CD239 is entirely expressed in adult mouse kidneys. Because the short-tail isoform, termed BCAM, was discovered as a tumor antigen in ovarian carcinoma (16), it may appear in renal carcinomas. In the inner stripe and inner medulla, the cytoplasm of the renal tubules and blood vessels was stained with the cyCD239 antibody (cyLu), indicating the presence of cytoplasmic domain fragments in the cells. Because the staining completely disappeared in the inner stripe and inner medulla lacking CD239, it was not a nonspecific reaction of the CD239 antibody (Fig. 4a). The cytoplasmic region of the long-tail isoform carries a SH3-binding motif, a spectrin-binding motif, a dileucine motif, and potential phosphorylation sites (36–38). Although further experiments are required, this fragment may function as a modulator of signal transduction and the cytoskeleton.

In adult mouse nephrons, CD239 is expressed in the epithelial cells of the glomeruli, Henle’s loop, distal tubules, collecting ducts, and the kidney capsule. In the vascular system, CD239 was observed in the renal and segmental arteries but not in the capillaries. CD239 co-localizes with laminin α5, that is a receptor ligand. CD239 expression is dramatically reduced in various tissues of mouse embryos lacking laminin α5 chain (26, 39). In contrast, it is significantly increased in the heart of transgenic mice overexpressing laminin α5 chain (26). As described in previous studies, CD239 is expressed in laminin α5-dependent manner in vivo. However, despite laminin α5 expression, CD239 was not observed in the proximal tubule and capillary. The cytoplasmic domain of CD239 binds to erythrocyte and non-erythrocyte spectrins and modulates networks in the cytoplasm (37, 38, 40, 41). Proximal tubule and capillary endothelial cells form highly organized intracellular spectral networks (42–44). This finding indicates that a distinct pathway modulates spectrin networks without anchoring CD239. In CD239-null mice, a distinct pathway may substantially compensates for the impairment of spectrin networks in various tissues.

The localization of UMOD was disrupted in distal tubules lacking CD239. Disturbances in UMOD localization are associated with CKD, kidney stone formation, and hypertension (30). However, as no alteration in renal function was observed in CD239-null mice, the mislocalization of UMOD appears to be due to a distinct signal in the epithelial cells of the distal tubule. Disrupted localization of the Na–K–Cl cotransporter (NKCC2) was also observed in CD239-null kidneys. Reportedly, NKCC2 is well co-localized with spectrin βII in the distal tubules (42). Although the interactions between NKCC2 and spectrins require further investigation, it is likely that the spectrin network, which lacks CD239 anchoring, influences the localization of this transport protein.

Although the CD239-null mice were healthy and developed normally, the stability of the distal renal tubules deteriorated. An experiment using an adenine-feeding model indicated that distal tubules lacking CD239 exhibited more extensive injury than in control mice fed with an adenine-supplemented diet. In humans, null phenotype red blood cells, Lu null or Lu(a–b–), which lack Lutheran system antigens, have been identified using traditional agglutination techniques (45). Pedigree analysis has further shown that Lu null arises from any of three genetic backgrounds: recessive, dominant, and X-linked (15). Among the inactivating mutations, the recessive type is the only true Lu-null phenotype arising from homozygosity for a silent allele at the gene locus, indicating that individuals represent natural human CD239 knockouts. Similar to the CD239-null mice, these individuals exhibited no obvious associated pathologies. Although it is unclear whether the stability of the distal renal tubule deteriorates in individuals, the kidney should be injured under loading conditions that do not normally lead to CKD. In(Lu) is a rare dominant suppressor of Lutheran blood group antigens. In(Lu) results from heterozygous mutations in KLF1 in the presence of a normal KLF1 allele (46). KLF1 is a transcription factor required for terminal differentiation of erythroid cells. Therefore, Lutheran blood group antigens seem to be expressed in the kidney, carrying the genetic background of In(Lu). The Lu null of the X-linked type results from the recessive X-borne inhibitor gene, *XS2* (47). Although it is necessary to examine the expression of Lutheran blood group antigens in the kidneys, it is likely that this type inhibits or suppresses these molecules throughout the body. Our results using a mouse model indicated that the CD239-null status is a predisposing factor for CKD. Lu-null individuals may need to avoid diets that impose a high renal load.

## METHODS

### Antibodies and regents

The primary antibodies used in this study are listed in Supplemental Table 1S (25, 48). Adenine was purchased from Merck (Kenilworth, NJ).

### Animals

C57BL/B6 mice, aged 8–9 weeks, were purchased from Oriental Yeast Co., Ltd. (Tokyo, Japan). *Bcam* knockout and CD239-null mice were generated as described in our previous studies (25). All animals were maintained in a standard Specific-Pathogen-Free (SPF) room at the institutional animal facility. Experiments were performed in accordance with the animal care guidelines of the Ehime University School of Medicine and Tokyo University School of Pharmacy and Life Science.

### Immunohistochemistry

The mouse kidneys were cut in the transverse plane. Oriented tissues were frozen in Tissue-Tek Optimum Cutting Temperature Compound (Sakura Finetek). Sections (6 µm) were cut using a cryostat and air-dried. After blocking with 10% normal goat serum, the sections were incubated with primary antibodies. Bound rabbit and rat IgG were detected using secondary antibodies conjugated to Alexa Fluor 594 and Alexa Fluor 647, respectively (Thermo Fisher Scientific). The signals of Alexa Fluor 647 are converted to green in the figures. After washing with phosphate-buffered saline (PBS) (-), the sections were mounted in 90% glycerol containing 0.1 × PBS and 1 mg/mL *p*-phenylenediamine. To generate images of the entire kidney section, gradually shifted images were captured at 10x magnification using a BZ-X810 microscope (Keyence, Osaka, Japan). A series of files were combined into one image of the whole kidney section using a BZ-X800 analyzer.

### Adenine-Induced Renal Tubule Injury

The mouse CKD model was generated by adenine feeding with some modifications (33). Wild-type and CD239 knockout mice were maintained on a control diet (CL-2; CLEA Japan, Tokyo, Japan). When the mice were 10 weeks old, feeding was switched to a 0.2% adenine-supplemented diet with free access to distilled water for 4 weeks. After adenine feeding, the mice were fed a control diet for 1 week and then blood samples were collected under isoflurane anesthesia and sacrificed by cervical dislocation. For histological analysis, mice were dissected after 2 weeks of adenine feeding.

### Blood Biochemical Analysis

The collected blood samples were centrifuged at 3,000 x g (4°C) and plasma was separated. To evaluate kidney function, plasma was subjected to animal blood biochemical analysis (Oriental Yeast, Tokyo, Japan).

### Statistical Analysis

Each bar represents the mean of three assays. The error bars indicate the standard deviation. Statistical significance was determined using Student’s *t* test.

### Study Approval

Animal studies using C57BL/6 mice were approved by the Tokyo University School of Pharmacy and Life Science Committee on the Care and Use of Laboratory Animals (P25-06). All animal experiments were approved by the Animal Care and Use Committee of the Ehime University Graduate School of Medicine (05-O-70-1.16).

## Author Contributions

Y.K. designed the research, conducted experiments, analyzed data, and wrote the manuscript. K.H., Y.Y and J.I conducted experiments, and analyzed data. M.T. and T.S. provided resources for the experiments. M.K. designed the research, analyzed data, and edited the manuscript. All authors discussed the results and provided comments on the manuscript.

## Acknowledgments

We would like to thank Dr Msumi Matsunuma of the Laboratory of Cellular Biochemistry for her assistance and helpful discussion. We also thank Ms. Harumi Kanno and Ms. Mika Kaneta in the Kanagawa laboratory for their technical assistance. We further thank the Division of Medical Research Support at the Advanced Research Support Center (ADRES), Ehime University, for technical support with tissue staining (Imaging Analysis Support Facility) and for maintaining the welfare of the animals used in this study (Laboratory Animal Facility).

## Competing interests

The authors declare no competing financial and non-financial interests.

## Funding

This work was supported in part by grants to YK (20K07622 and 23K07728) from the JSPS KAKENHI Grant, Japan and to YK (JPJSBP120248814) from the JSPS Bilateral Joint Research Projects, Japan.

## Notes

### Competing Interest Statement

The authors have declared no competing interest.

### Summary of Updates

Title was changed, and the revised manuscript was uploaded.

