## Supplemental Figures and Table for "CD239 deficiency in distal tubules impairs cell polarity and increases susceptibility to renal injury"

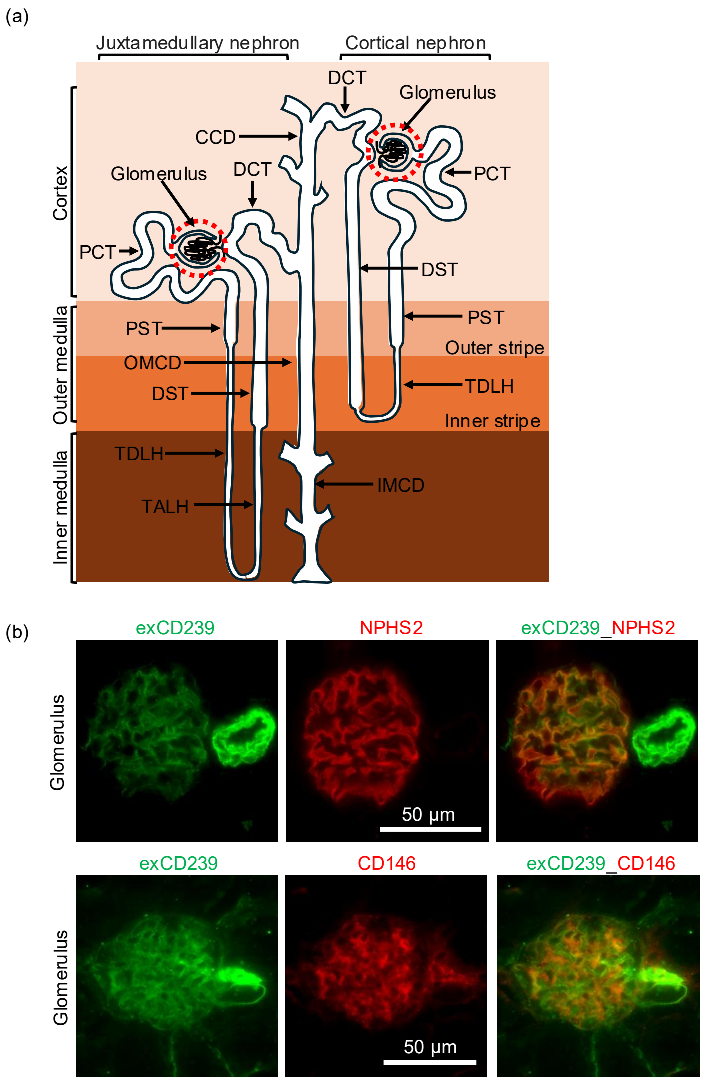


**Figure 1S. Expression of CD239 in glomeruli.**
(a) Schematic of the juxtamedullary and cortical nephrons. Glomeruli are surrounded by dotted circles. (b) Immunostaining of mouse kidney section. The sections were doubly stained with antibodies against exCD239 (green) and NPHS2 (red) or CD146 (red), respectively. The images were captured at high magnification. Bar: 50 µm.

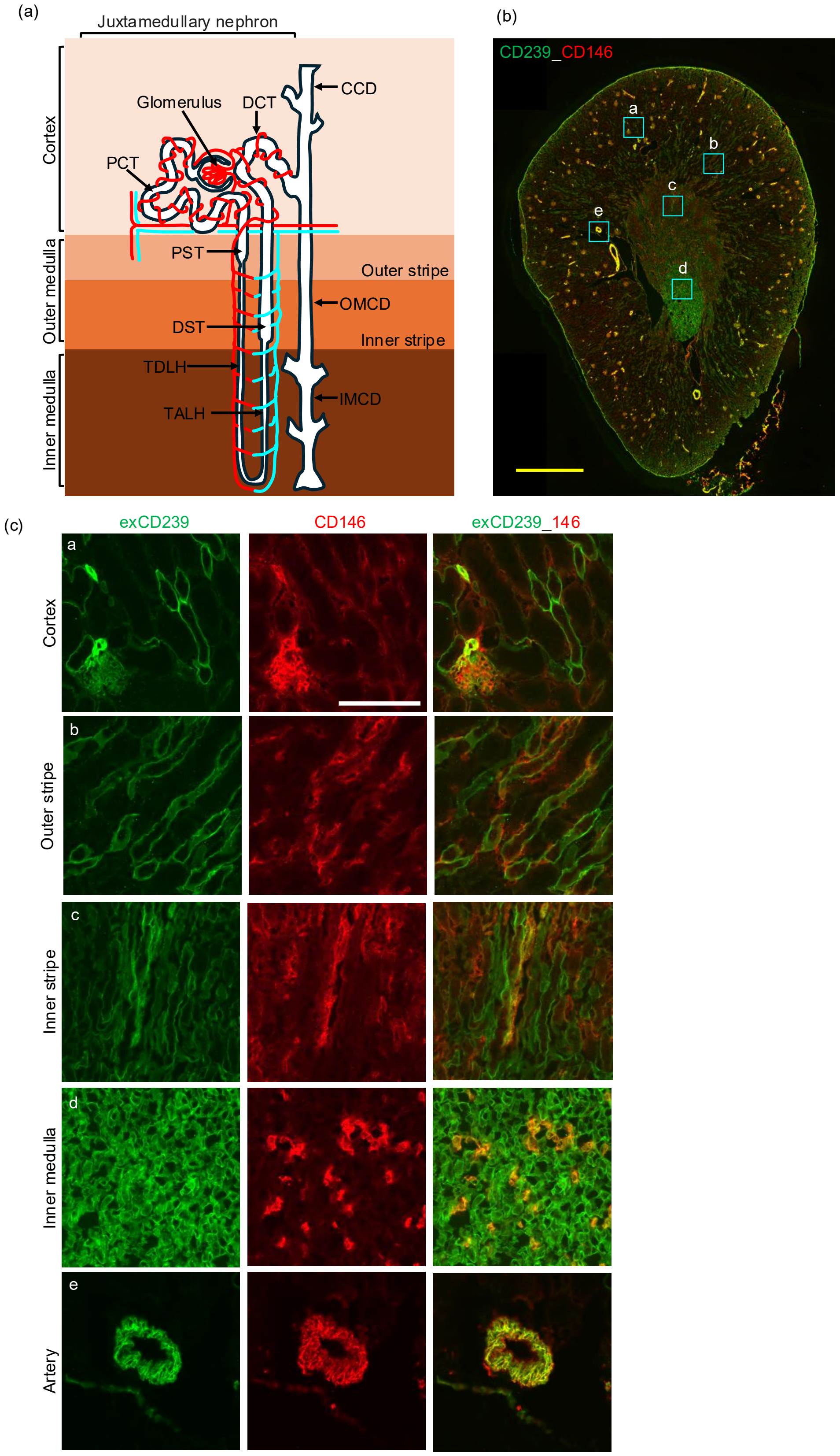


**Figure 2S. Expression of CD239 in blood vessels.**
(a) Schematic of nephron and vascular system. Renal arterioles and venules are depicted by red and light blue lines, respectively. (b) Immunostaining of a whole mouse kidney section. The sections were doubly stained with antibodies against exCD239 (green) and CD146 (red), respectively. Bar: 1.0 mm. (c) High magnification images in each anatomical region. The locations are indicated by blue squares on the image of the whole kidney section. Bar: 100 µm.

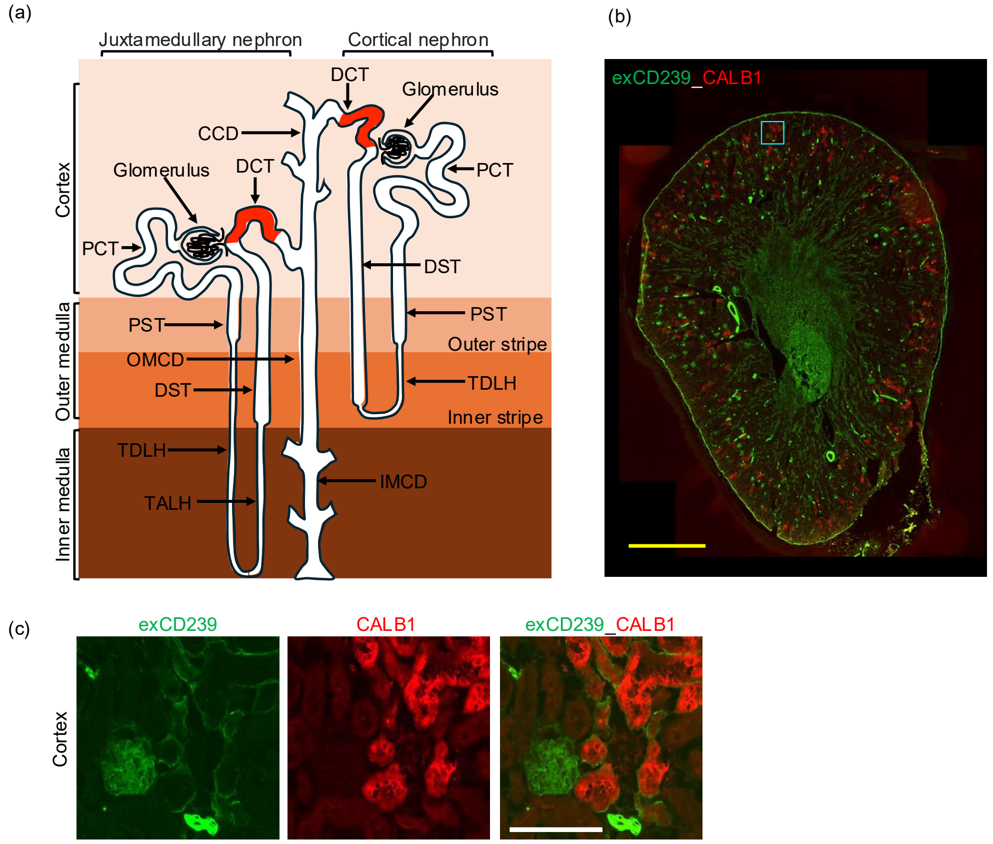


**Figure 3S. Expression of CD239 in distal convolved tubule (DCT).**

(a) Schematic of juxtamedullary and cortical nephrons. DCT is defined with localization of CALB1 (red). (b) Immunostaining of a whole mouse kidney section. The sections were doubly stained with antibodies against exCD239 (green) and CALB1 (red), respectively. Bar: 1.0 mm. (c) High magnification images in each anatomical region. The location is indicated by a blue square on the image of the whole kidney section. Bar: 100 μm.

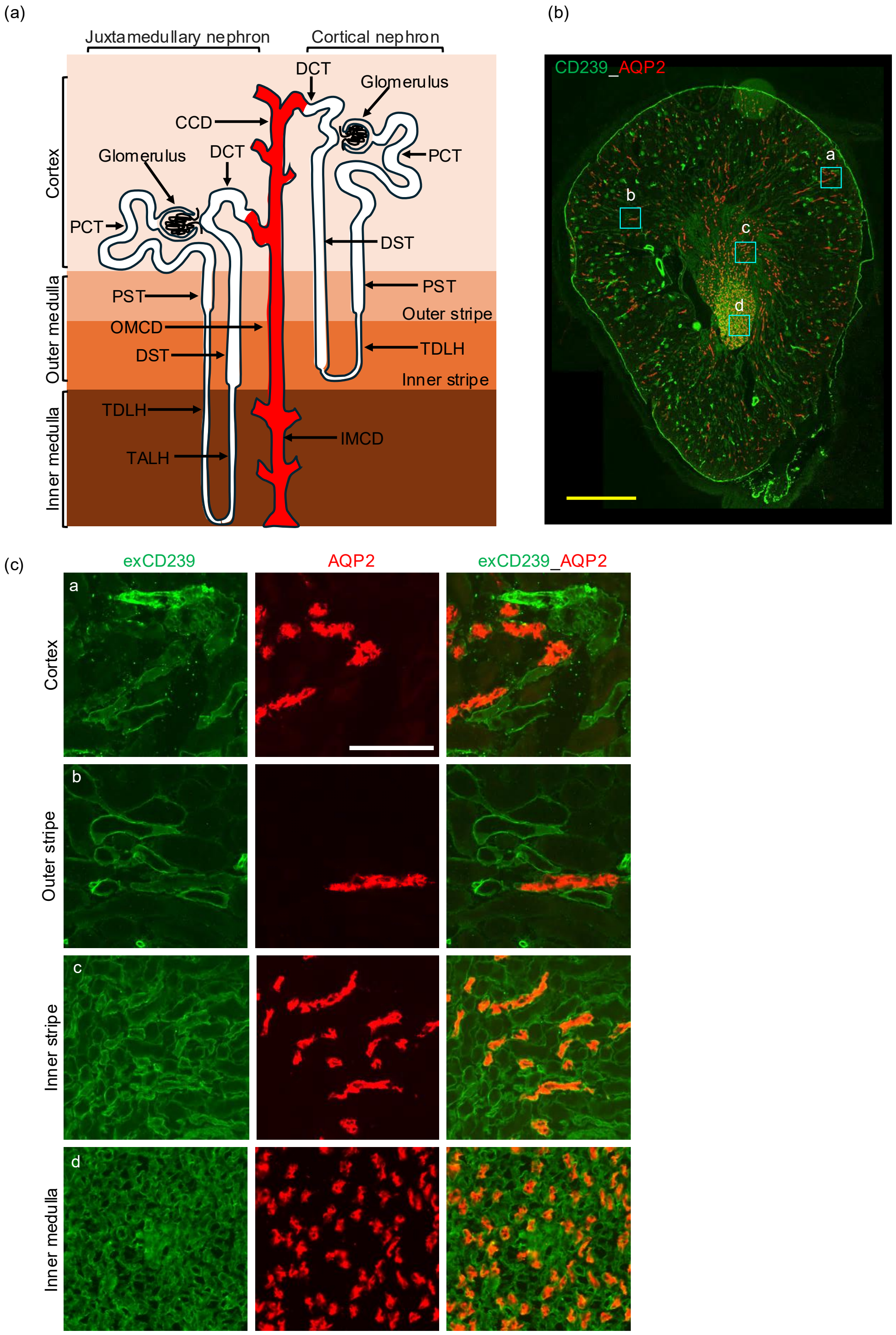


**Figure 4S. Expression of CD239 in collecting ducts (CD).**
(a) Schematic of juxtamedullary and cortical nephrons. CD are defined with localization of AQP2 (red). (b) Immunostaining of a whole mouse kidney section. The sections were doubly stained with antibodies against exCD239 (green) and AQP2 (red), respectively. Bar: 1.0 mm. (c) High magnification images in each anatomical region. The locations are indicated by blue squares on the image of the whole kidney section. Bar: 100 μm.

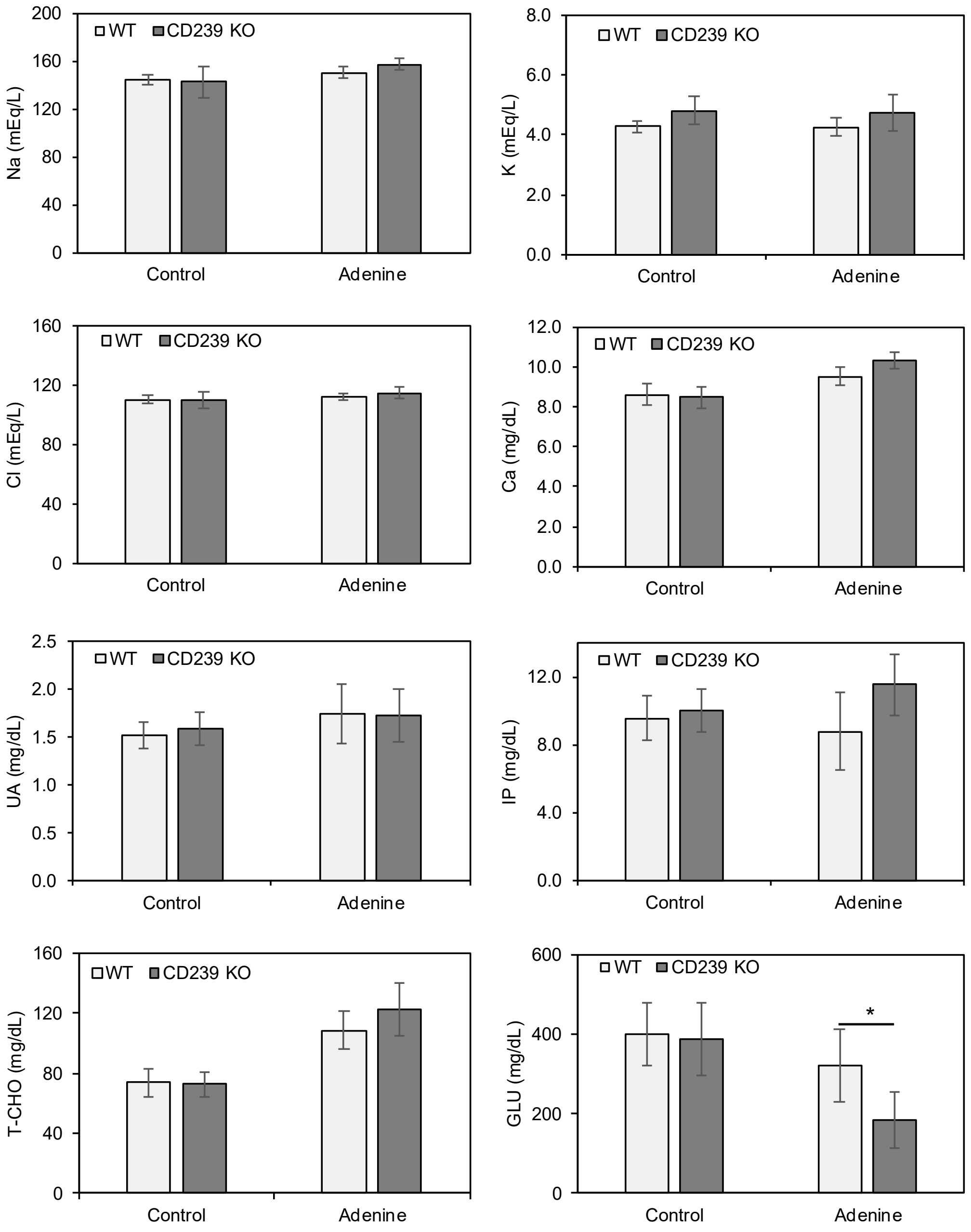


**Figure 5S. Blood biochemical analysis in the adenine-induced kidney injury mouse model.**
Influence of adenine supplementation on serum markers of kidney function. After feeding the adenine-supplemented diet, blood was collected from the wild-type and CD239-null mice. Serum samples were analyzed for Na, K, CL, Ca, uric acid (UA), inorganic phosphorus (IP), total cholesterol (T-CHO) and glucose (GLU). *, $p < 0.05$ by Student's t-test.

| Table 1S. Primary antibodies | | | | | |
|---|---|---|---|---|---|
| Antibody to | Epitope | Clone /ab no. | Host/Antigen Species | subclass | Source/Reference |
| CD239 | Extracellular domain | 10-5 | Rat/Mouse | $IgG_{2a}$ | Miura et al., eLife, 2018 |
| CD239 | Intracellular domain | cyLu | Rabbit/Mouse | - | Gift from Dr. Miner, Washington University in St.Louis |
| APQ1 | C-terminus | 20101 | Rabbit/Human | - | BiCell Scientific, Maryland Heights, MO |
| APQ2 | C-terminus | 20101 | Rabbit/Human | - | BiCell Scientific, Maryland Heights, MO |
| UMOD | - | 774056 | Rat/Mouse | $IgG_{2a}$ | R&D Systems, Minneapolis, MN |
| CALB1 | N-terminus | D1I4Q | Rabbit/Human | - | BiCell Scientific, Maryland Heights, MO |
| NKCC2 | N-terminus | 20301 | Rabbit/Mouse | - | BiCell Scientific, Maryland Heights, MO |
| LAMC1 | LN and LEa domains | 1083+ | Rabbit/Mouse | - | Garbe et al., Biochem J, 2002 (48) |
| F4/80 | - | BM8 | Rat/Mouse | $IgG_{2a}$ | BioLegend, San diego, CA |
| CD146 | Extracellular domain | m146 pAb | Rabbit/Mouse | - | In this study |
| NPHS2 | C-terminus | | Rabbit/Human | - | Abcam, Cambridge, MA |
| NID1 | - | ELM1 | Rat/Mouse | $IgG_{2a}$ | Merck, Kenilworth, NJ |
| BSND | C-terminus | 20402 | Rabbit/Mouse | | BiCell Scientific, Maryland Heights, MO |
